# Cigarette smoke extract and palmitic acid exposed patient-derived iPS-RPE cells as a model for AMD

**DOI:** 10.64898/2026.10.30.763204

**Authors:** Rabina Kumpakha, Mohd. Akbar Bhat, Ganesh Satyanarayana, Nady Golestaneh, Sayantan Datta

## Abstract

**Purpose:** Age-related macular degeneration (AMD) is a multifactorial disease in which dysfunction and loss of retinal pigment epithelial (RPE) cells contribute to disease progression. Besides genetics, cigarette smoking and high fat diet are important environmental and metabolic risk factors for AMD and are associated with RPE dysfunction. However, the effects of combined environmental and metabolic stress on AMD-derived RPE cells remain poorly understood. In this study, we used patient-derived induced pluripotent stem cell-derived RPE (iPSC-RPE) cells to examine the effects of cigarette smoke extract (CSE) and palmitic acid (PA) on RPE cell function.

**Methods:** iPSCs derived from non-AMD and AMD donors were differentiated into RPE cells and characterized based on morphology, RPE-specific marker expression, polarity, gene expression, and barrier function. Mature iPSC-RPE cells were exposed to different concentrations of CSE and PA to determine experimental conditions. Cell death, epithelial-to-mesenchymal transition (EMT)-like morphological changes, stress-related gene expression, RPE polarity, TEER, VEGF-A and PEDF secretion, and intracellular lipid accumulation were then evaluated following CSE, PA, or combined CSE and PA exposure.

**Results:** Differentiated iPSC-RPE cells developed typical pigmented, hexagonal morphology, expressed RPE-specific markers, and formed polarized monolayers. Both CSE and PA increased cell death in a dose-dependent manner in the control and AMD-iPSC-RPE, with AMD-iPSC-RPE exhibiting greater susceptibility, particularly those from patients with dry AMD. CSE and PA also induced also induced EMT-like morphological changes and increased the expression of *ZEB1*, *NQO1*, and *IL8*. Furthermore, CSE and PA disrupted the polarized distribution of ZO-1 and Best1, significantly reduced the polarity ratio, decreased PEDF and VEGF-A secretion and increased intracellular lipid accumulation. Combined CSE and PA treatment produced more pronounced effects than either treatment alone in AMD lines.

**Conclusions:** AMD-iPSC-RPE cells, particularly those derived from dry AMD donors, exhibit increased susceptibility to environmental and metabolic stress. Exposure to CSE and PA revealed this vulnerability through alterations in cell survival, stress responses, epithelial morphology, polarity, secretory function, and intracellular lipid accumulation. These findings highlight the effects of CSE and lipid stress on RPE metabolism and underscore the utility of patient-derived iPSC-RPE cells as a model for investigating how environmental and metabolic factors contribute to RPE dysfunction and AMD pathogenesis.

## Introduction

Age-related Macular Degeneration (AMD) is a multifactorial degenerative disease of the macula and one of the leading causes of irreversible vision loss in individuals over 55 years of age. Approximately 196 million people were estimated to be affected by AMD globally in 2020 and this number is projected to increase to 288 million by 2040 [1]. AMD involves progressive dysfunction and degeneration of several components of the outer retina, including photoreceptors, retinal pigment epithelium (RPE), Bruch’s membrane, and choriocapillaris [2]. Although AMD is traditionally classified into neovascular (wet AMD) and non-neovascular (dry AMD) forms, dysfunction and degeneration of the RPE are important features of both early and advanced stages of the disease. Dry AMD can be further classified into early, intermediate, and advanced stages, with the advanced stage characterized by geographic atrophy (GA). Early and intermediate dry AMD are the most common forms and affect millions of individuals worldwide. Progressive disease can lead to RPE dysfunction, degeneration, and the development of areas of atrophy that ultimately progress to GA, resulting in irreversible loss of central vision. Although effective treatments are available for wet AMD, therapeutic options for dry AMD, particularly advanced or late stage AMD remain limited [3]. Consequently, understanding the cellular and molecular mechanisms underlying RPE dysfunction is critical for developing effective therapeutic strategies for dry AMD.

Early pathological changes in dry AMD are closely associated with cellular stress and dysfunction of the RPE, which can be influenced by environmental, metabolic, genetic and lifestyle-related factors. The RPE is a highly specialized monolayer of polarized cells that performs several essential functions required for retinal homeostasis. These include phagocytosis and degradation of photoreceptor outer segments, support of visual cycle, transport of oxygen, nutrients and metabolic waste between the retina and choroid, and maintenance of the outer blood-retinal barrier [4]. Because of its central role in maintaining the retinal microenvironment, disruption of RPE function can, therefore, impair multiple aspects of retinal homeostasis. Hence, persistent RPE dysfunction and cellular stress can consequently contribute to photoreceptor degeneration and the progression of AMD, particularly toward advanced forms.

The development of AMD is influenced by a combination of aging, genetic susceptibility, and environmental factors. Stressors such as smoking and high fat diet are known to be strong risk factors that contribute to AMD [5–8]. Smoking has shown association with both geographic atrophy and neovascular AMD. Cigarette smoke exposure induces oxidative stress responses, promotes production of reactive oxygen species (ROS) and causes lipid metabolism disorder leading to RPE dysfunction [9,10]. Oxidative stress can also interact with inflammatory pathways, potentially contributing to progressive RPE dysfunction [11].

In addition to oxidative stress, lipid metabolism dysregulation is an important component of AMD pathobiology. Aging and AMD are associated with changes in lipid composition and accumulation of lipid-rich material in the RPE-Bruch’s membrane complex [8,12]. Recent studies have also highlighted interactions between altered lipid metabolism and complement activation in AMD, suggesting that lipid abnormalities may contribute to chronic inflammatory responses in the outer retina [13]. Palmitic acid (PA), a common saturated fatty acid, can produce cellular stress when present in excess. PA has been found to induce injury in retinal epithelial cells and promote inflammatory response [14,15]. Increased exposure to saturated fatty acids like PA not only can alter lipid storage and metabolism but also causes drop-in cellular metabolic activity increasing ROS generation and promote cell death due to lipotoxicity in susceptible cells [5].

Environmental and metabolic stressors may act synergistically in AMD. Cigarette smoke can increase oxidative stress and alter cellular metabolism, while abnormal lipid accumulation may further increase cellular stress and inflammatory signaling. It is therefore possible that exposure to cigarette smoke and excess saturated fatty acids places an additional burden on RPE cells that are already susceptible because of disease-associated genetic or cellular characteristics. Understanding how these stressors interact may provide insight into why some individuals develop more severe RPE dysfunction than others.

Induced pluripotent stem cell (iPSC) technology has been widely used to investigate disease-associated phenotypes and cellular responses in AMD and other retinal diseases [16–18]. Patient-derived iPSCs can be differentiated into RPE cells that develop characteristic morphological, molecular, and functional properties of RPE, while retaining the donor’s genetic background. Patient-derived iPSC-RPE models provide an opportunity to compare responses between individuals with and without disease under controlled experimental conditions.

In the present study, we used iPSC-derived RPE cells from age matched non-AMD and AMD donors to investigate the effects of cigarette smoke extract (CSE) and PA on RPE survival and function. We first characterized the differentiated cells based on their morphology, RPE-specific marker expression, polarization, gene expression, and barrier function. We then examined the effects of CSE and PA, individually and in combination, on cell death, EMT-like changes, cellular stress responses, RPE polarity, VEGF-A and PEDF secretion, and intracellular lipid accumulation. We hypothesized that AMD-derived iPSC-RPE cells would be more susceptible to environmental and metabolic stress than non-AMD iPSC-RPE cells, and that combined exposure to CSE and PA would produce more pronounced cellular dysfunction than either stressor alone in which cells, do you mean in both control and AMD-iPSC-RPE?.

## Results

### Patient demographics and genotyping of cell lines

The iPSC-RPE lines used in this study represented a reference line (IMR90-4), a non-AMD donor with a family history of AMD, and donor-derived lines from individuals with dry AMD and Stage 4 advanced dry AMD (Adv AMD). The donor-derived AMD lines were obtained from aged individuals, including an 86-year-old male with dry AMD and an 80-year-old female with Adv AMD. Genotyping of AMD-associated loci revealed differences in *CFH*, *ARMS2*, *C2*, and *C3* risk genotypes among the cell lines (supplementary data). Notably, both the dry AMD and Adv AMD lines carried a low-risk *CFH* genotype (TT), high-risk *ARMS2* genotype (TT), low-risk *C2* genotype (GT), and moderate-risk *C3* genotype (CG). The non-AMD donor with a family history of AMD also carried a low-risk CFH genotype (TT). Thus, the AMD-derived lines shared a similar genetic risk profile across the four loci examined, particularly with respect to low *CFH* risk. For experiments examining the effects of an external stressor, we selected cell lines carrying the low-risk *CFH* genotype (TT) to minimize potential contribution of *CFH*-associated genetic risk and provide a more consistent genetic background for evaluating cellular responses to the external stressor.

### Morphological and functional characterization of non-AMD- and AMD-derived iPSC-RPE cells

The iPSCs derived from non-AMD and AMD donors were differentiated into RPE cells for 10 weeks. Following differentiation, mature iPSC-RPE cells appeared as highly pigmented hexagonal cells, resembling the morphology of native RPE (Figure 1). Brightfield microscopy demonstrated comparable morphology across the non-AMD, AMD, and IMR90-4 iPSC-RPE cell lines, with all cell lines exhibiting the characteristic hexagonal RPE morphology (Figure 1A).

**Figure 1:**
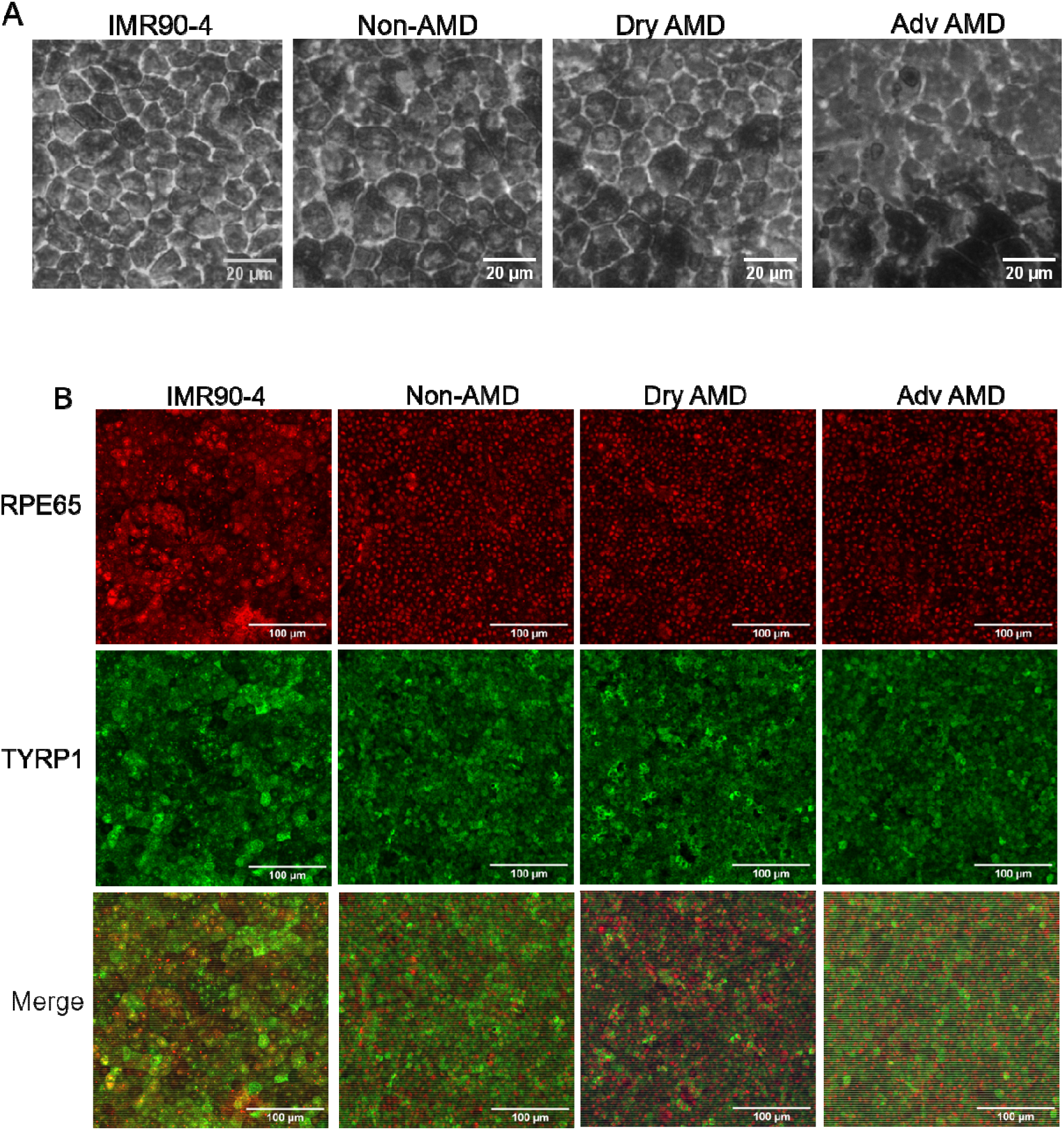
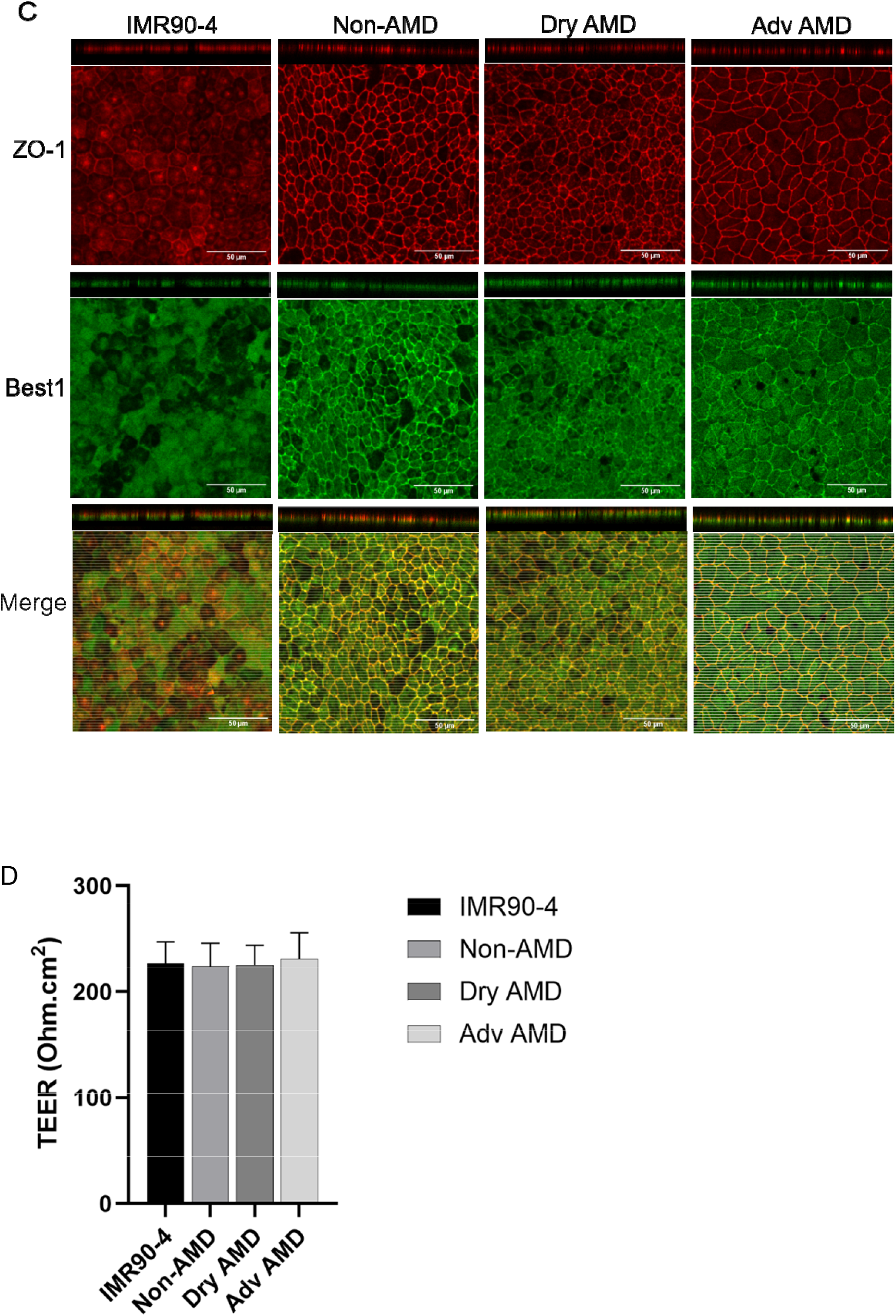
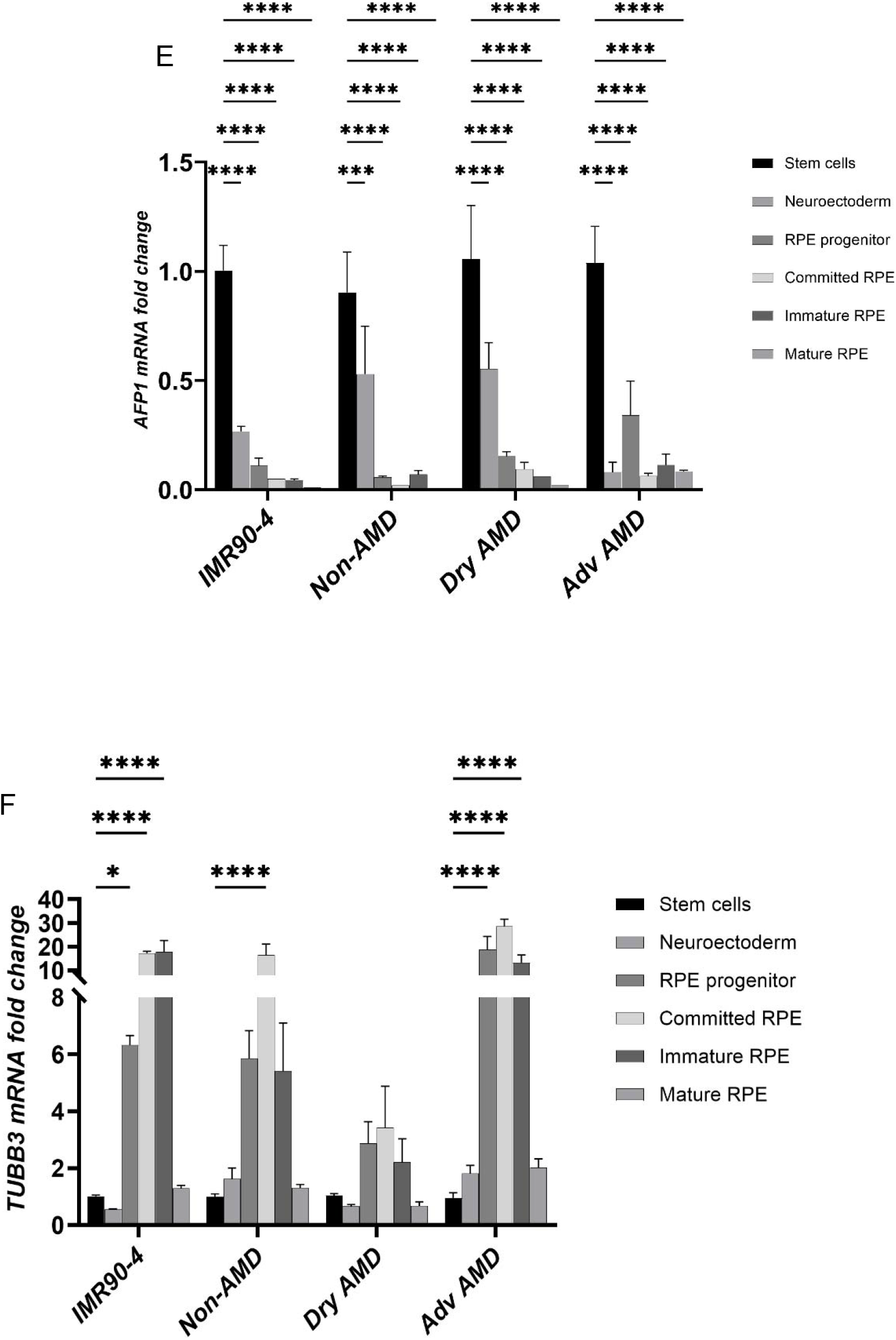
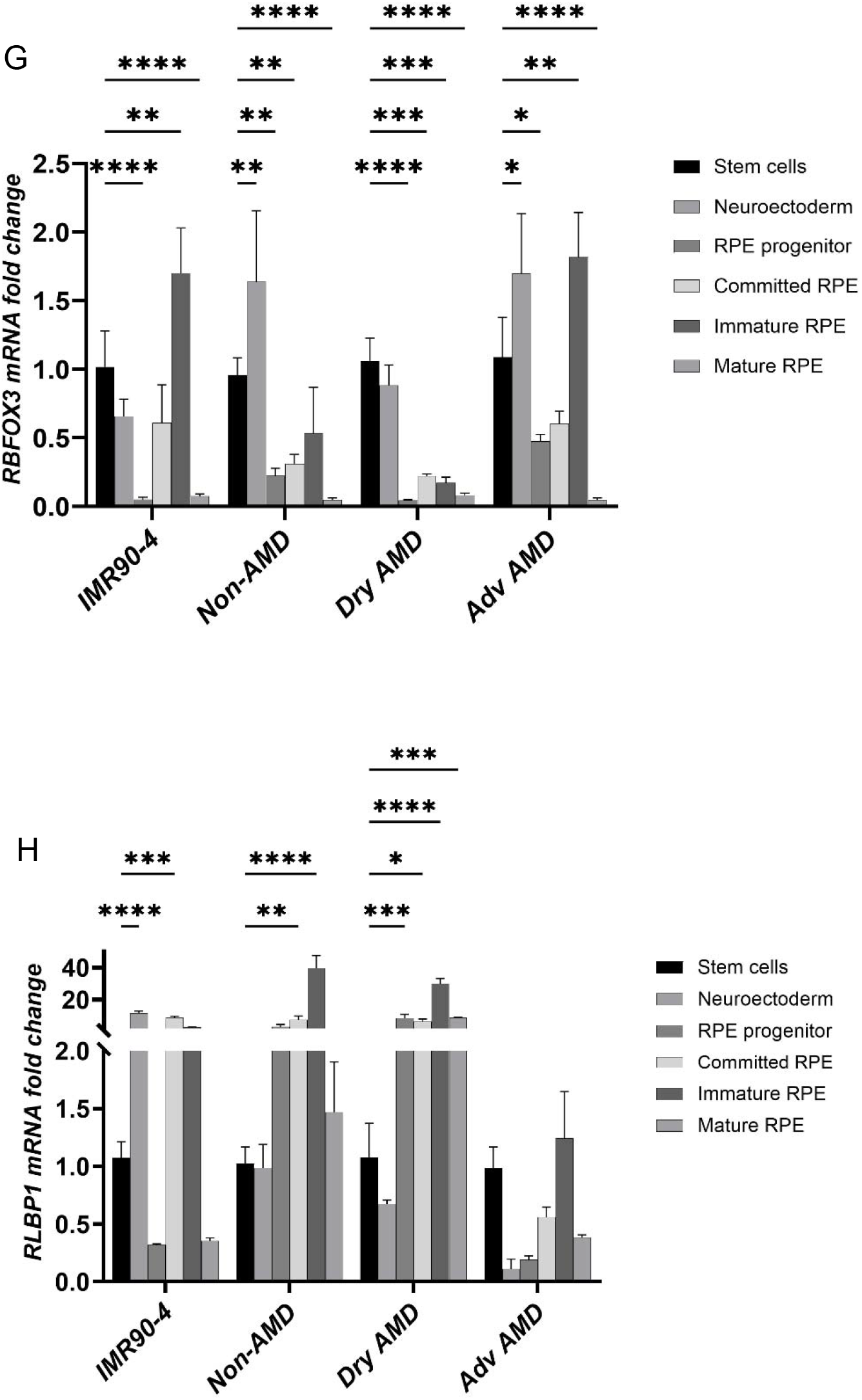
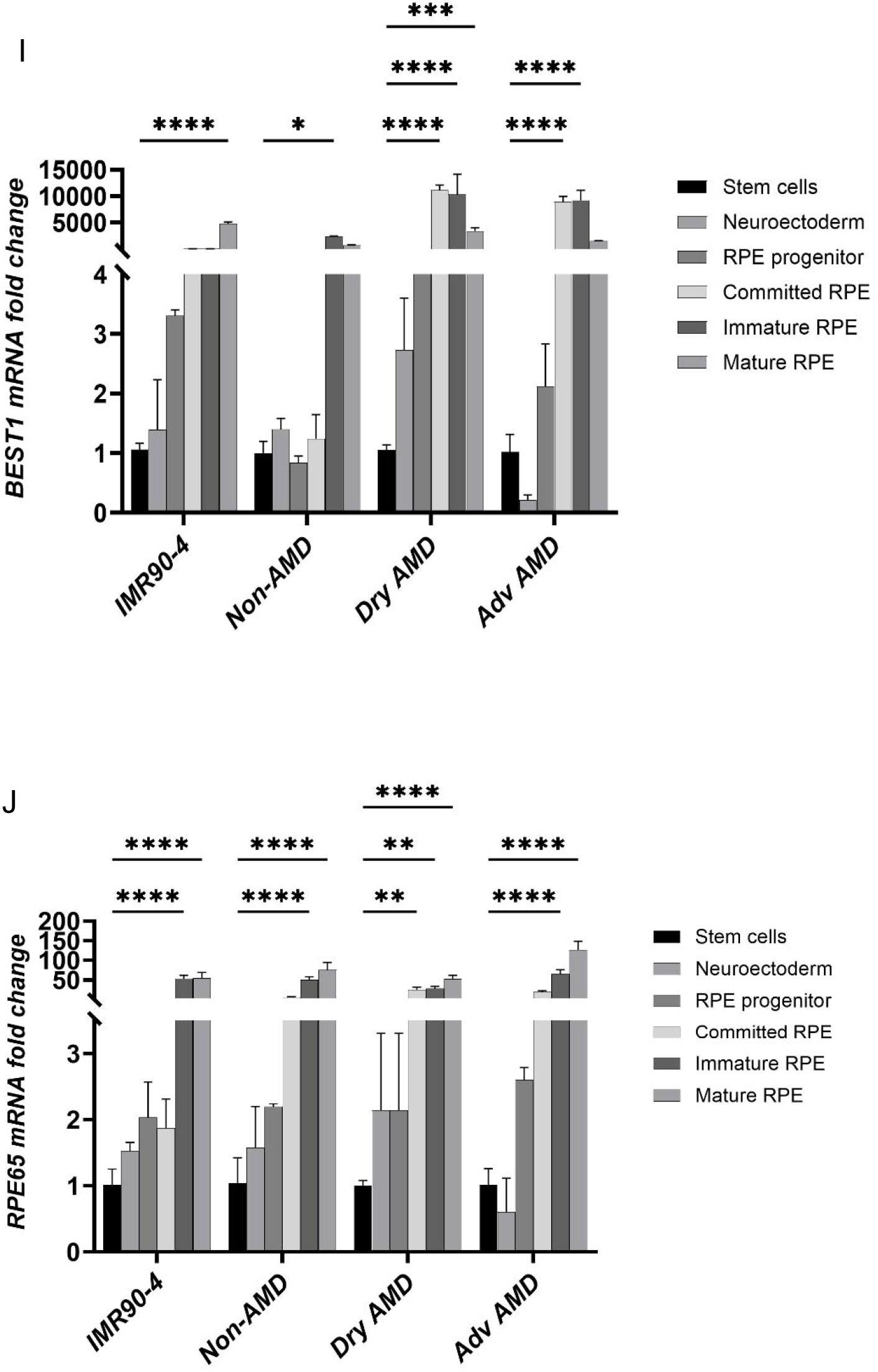
Morphology of mature iPSC-RPE cells derived from non-AMD, Dry AMD, and Adv AMD compared to mature IMR90-4 iPSC-RPE cells. (A) Brightfield images of mature all four iPSC cell lines showing characteristic hexagonal cellular structure and pigmentation. (B) Confocal images of mature iPSC-RPE cells immunostained with RPE specific markers; RPE65 (red) and TYRP1 (green). (C) Mature iPSC-RPE cells grown in Transwell inserts stained with Best1 and tight junction protein, ZO-1 for polarity. Representative two-dimensional confocal images and orthogonal views showing polarized RPE cells. Scale bar: 100 microns. (D) Transepithelial resistance of cells grown in a Transwell system. (E-G) qPCR for the stem cell specific markers; *AFP1*, *TUBB3* and *RBFOX3*, at different stages of differentiation. (H-J) qPCR for RPE cell specific markers; *RLBP1*, *BEST1* and *RPE65* at different stages of differentiation. Each cell line was differentiated into at least 3 clonal expansions, and the experiments were done in triplicates on those clonal lines. The data are presented in graphs as mean ± standard deviation from three independent experiments.

RPE differentiation was further confirmed by immunostaining for the RPE-specific markers RPE65 and TYRP1. All three mature iPSC-RPE cell lines showed positive staining for both markers (Figure 1B). To assess polarization, cells were cultured on Transwell filter inserts for 8 weeks and stained for the tight junction protein ZO-1 and the RPE marker Best1. Orthogonal views of z-stack confocal images demonstrated polarized localization of these proteins, with ZO-1 predominantly localized toward the apical region and Best1 toward the basolateral region of the cells (Figure 1C).

Barrier function was assessed by measuring transepithelial electrical resistance (TEER). TEER values were above 200 Ω·cm² in all cell lines, with no significant differences observed between the groups (Figure 1D). Gene expression analysis further confirmed RPE maturation. Expression of the stem/progenitor-associated markers *AFP1* (Figure 1E) *TUBB3* (Figure 1F), and *RBFOX3* (Figure 1G) decreased following differentiation, whereas expression of RPE-associated genes, including *RPE65* (Figure 1I), *RLBP1* (Figure 1J), and *BEST1* (Figure 1K), increased in mature iPSC-RPE cells Together, these findings confirmed the development of mature, polarized iPSC-RPE cells with characteristics consistent with native RPE.

### CSE and PA induce dose-dependent cell death in iPSC-RPE cells

To determine suitable concentrations of CSE and PA for subsequent experiments, mature iPSC-RPE cells were exposed to increasing concentrations of PA or CSE. The mature iPS-RPE cells were preliminarily treated with 0, 100, 250 and 500 µg/ml of CSE or 0, 100, 200 and 400 µM PA for 24 hours. Cell death was assessed using Hoechst and ethidium homodimer staining. Untreated control cells showed minimal cell death, whereas both CSE and PA increased cell death in a dose-dependent manner in both non-AMD and AMD-iPSC-RPE (Figure 2).

**Figure 2:**
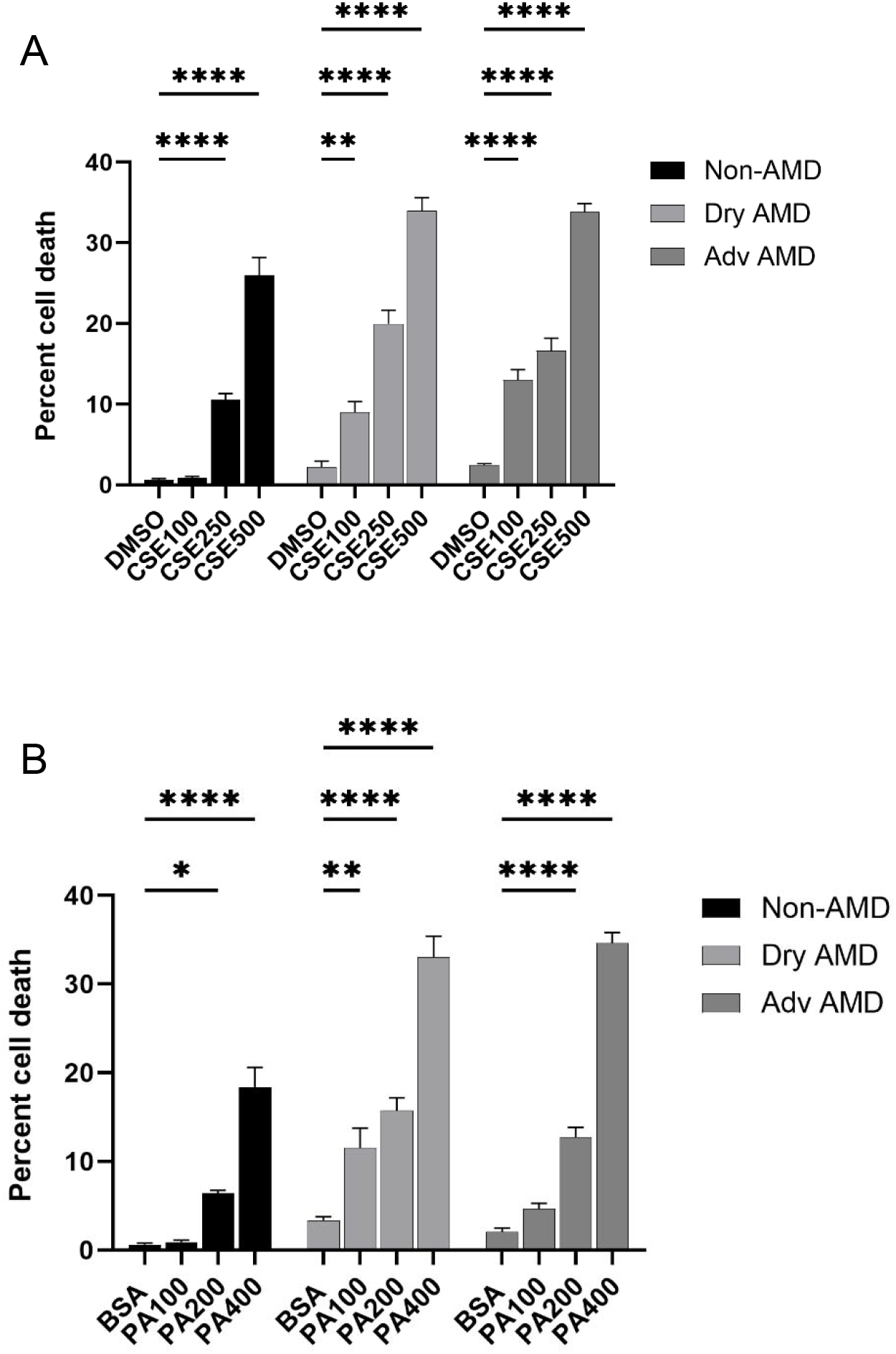
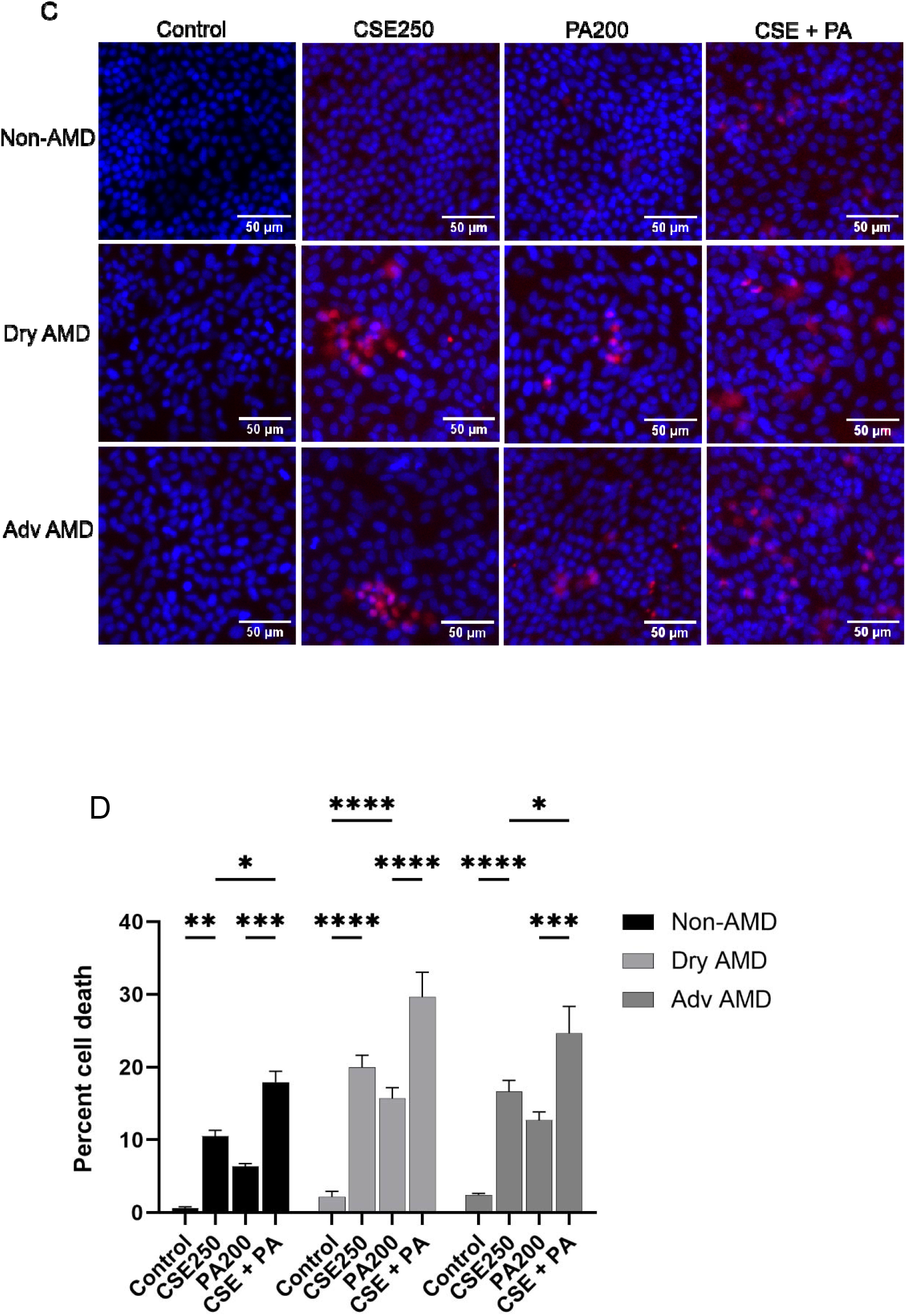
Effect of CSE and PA co-treatment on patient-derived mature iPS-RPE cells. Cells were assayed for cell death assay and EMT post treatments. (A) Quantification of dose dependent cell death assay for CSE treatment using Ethidium homodimer and Hoechst stain. (B) Quantification of dose dependent cell death assay for PA treatment using Ethidium homodimer and Hoechst stain. (C) Fluorescence images showing cell death with CSE and PA treatment. Blue stain indicates total cells, and red stain indicates dead cells. (D) Quantification of cell death assay from CSE and PA co-treatment. Each cell line was differentiated into at least 3 clonal expansions, and the experiments were done in triplicates on those clonal lines. Data shown in bar graphs are presented as mean ± standard deviation and from at least three independent experiments. The p-value of <0.05 is considered significant.

CSE exposure resulted in a concentration-dependent increase in cell death across all three cell lines. The highest concentration tested produced the significantly greater cell death in both non-AMD and AMD iPSC-RPE cells compared to their corresponding untreated control showing approximately 35% cell death in AMD cell lines and approximately 30% in non-AMD. Treatment with lower CSE concentrations resulted in approximately 20% cell death in both AMD iPSC-RPE cells compared to their control, while non-AMD showed approximately 10% cell death (Figure 2A).

A similar dose-dependent response was observed following PA exposure. The highest PA concentration resulted in the greatest cell death, whereas lower concentrations produced a comparatively smaller effect (Figure 2B). Overall, AMD-derived iPSC-RPE cells showed greater susceptibility to both CSE- and PA-induced cell death in dose dependent manner, compared with non-AMD cells.

Based on these dose-response experiments, 200 µM PA and 250 µg/mL CSE were selected for subsequent experiments. Mature iPSC-RPE cells were then treated with CSE, PA, or both treatments for 24 hours. Both individual treatments increased cell death compared with untreated controls as shown by ethidium bromide treatment and Hoechst staining, while combined CSE and PA exposure resulted in a further increase in cell death (Figure 2C and 2D). AMD-derived cells were more susceptible to the treatments than non-AMD cells. Among the AMD-derived cell lines, dry AMD cells showed the greatest sensitivity to the combined treatment (Figure 2D).

### CSE and PA induce EMT-like morphological changes and stress responses

To determine whether CSE and PA altered the epithelial phenotype of mature iPSC-RPE cells, cells were stained for the tight junction protein ZO-1 following 24-hour treatment. Untreated control cells maintained the characteristic epithelial morphology and organized ZO-1 staining. In contrast, exposure to CSE or PA resulted in morphological changes consistent with an EMT-like phenotype in all cell lines (Figure 3A).

**Figure 3:**
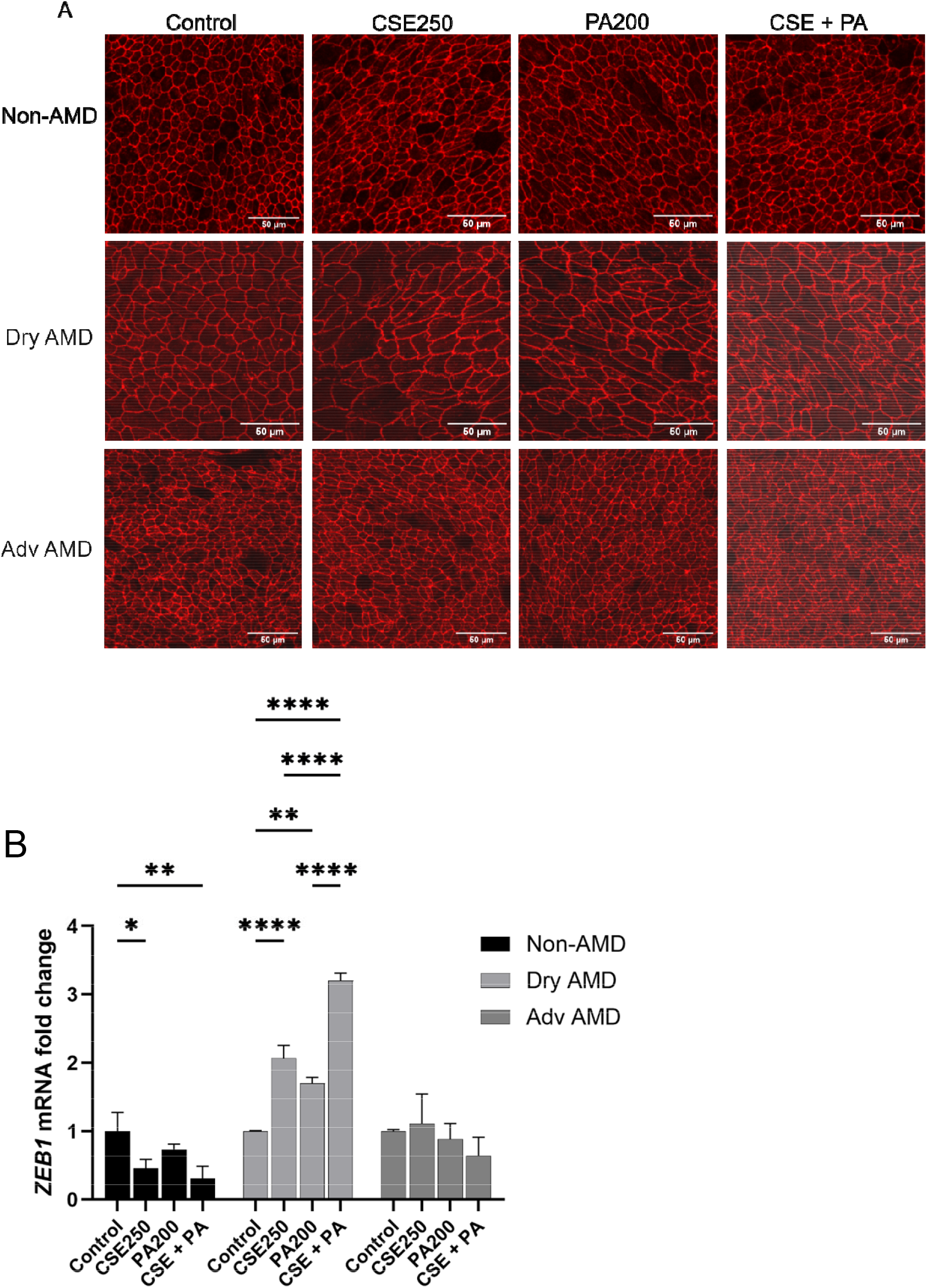
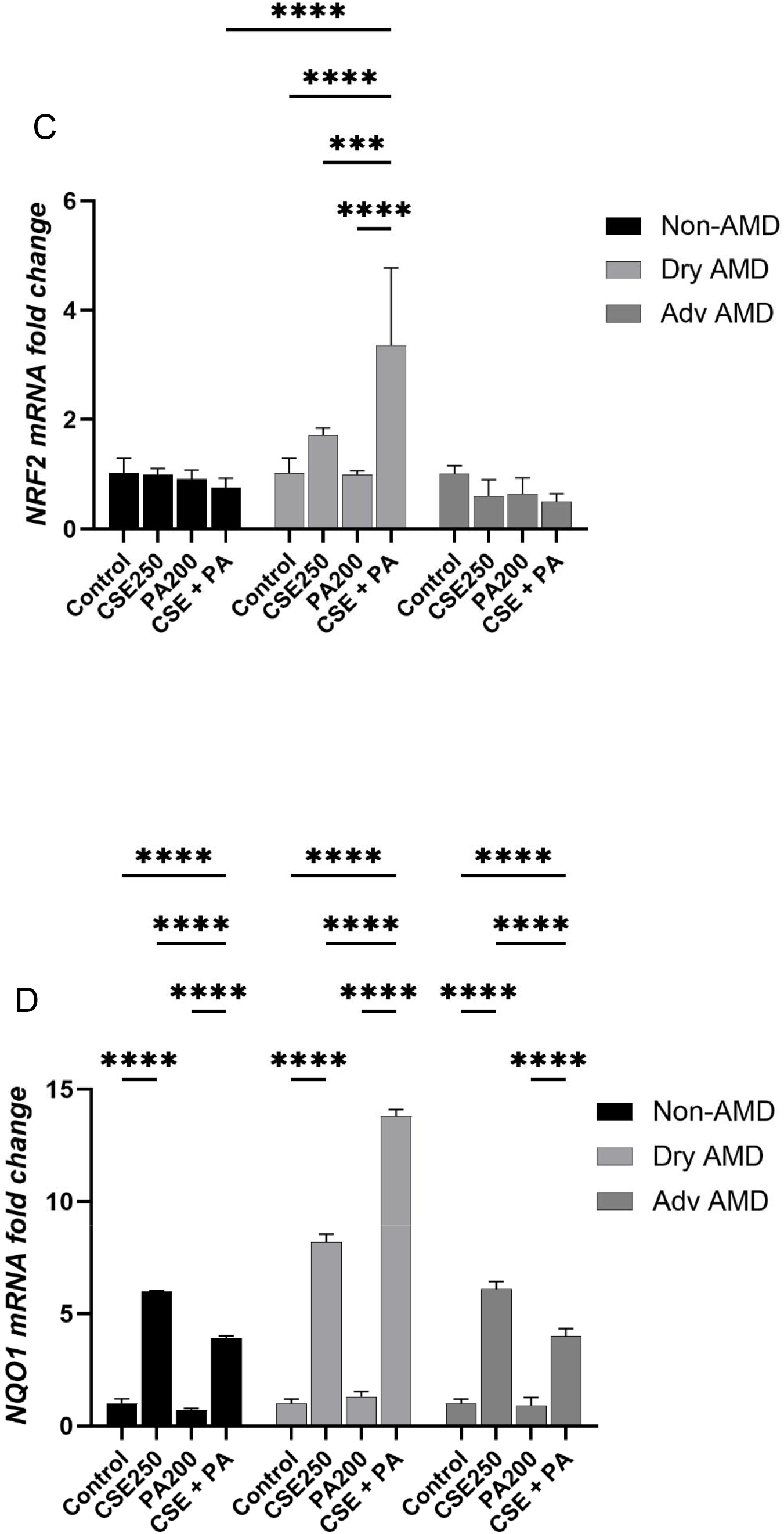
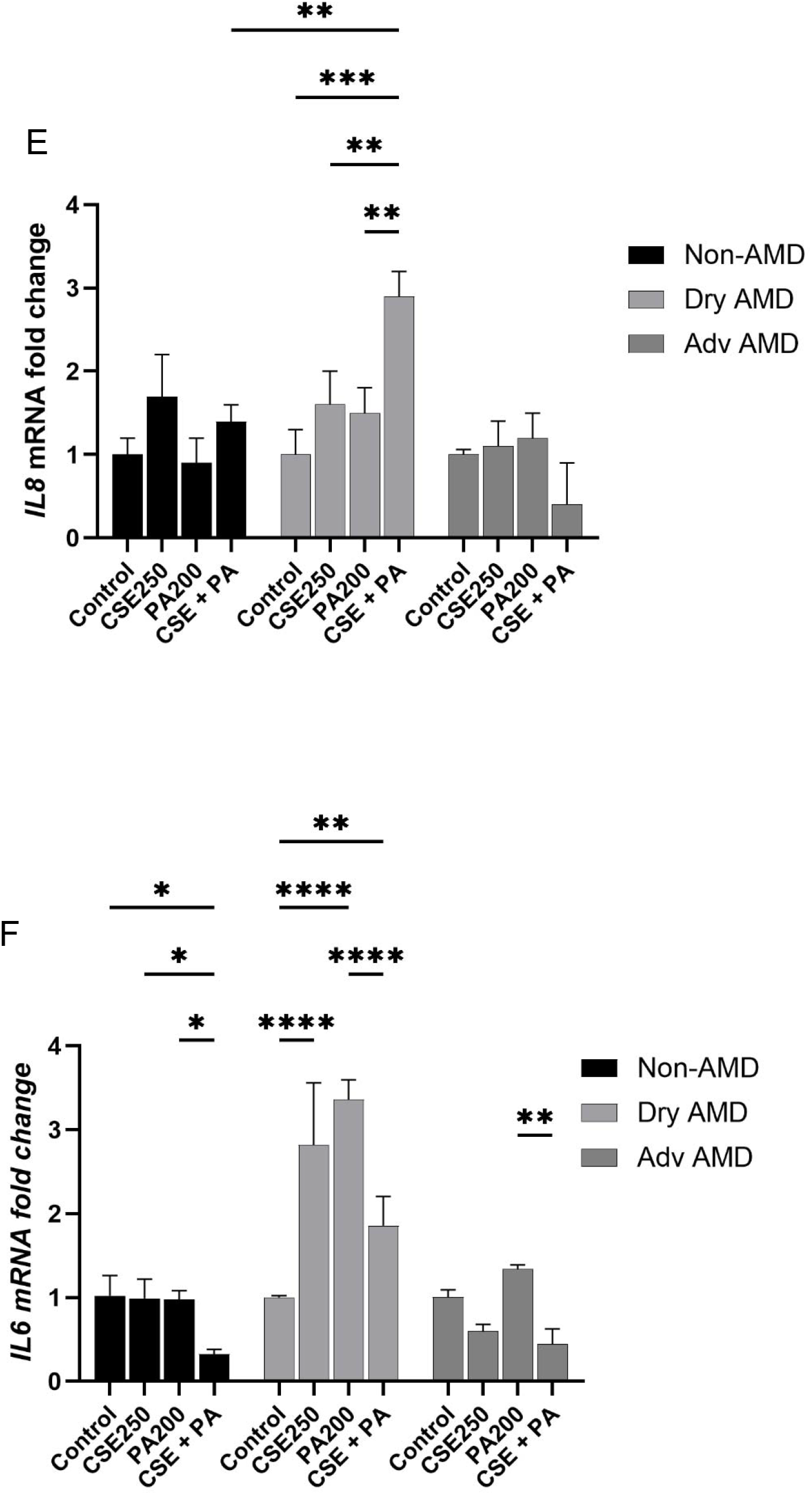
Stress response of human derived iPS cell lines post CSE and PA treatment. (A) Confocal images of mature non-AMD and AMD iPSC-derived RPE cells immunolabeled with tight junction protein ZO-1 showing strong EMT like morphology in CSE and PA treated cells. (B) Gene expression analysis using qPCR for EMT marker *ZEB1*, a master regulator for mesenchymal phenotype, represented as fold change. (C) Gene expression analysis using qPCR for *NQO1,* a transcription factor for downstream antioxidant genes. (D) Gene expression analysis using qPCR for *NRF2*, a master transcriptional regulator for antioxidants. (E) Gene expression analysis using qPCR for *IL8,* a primary pro-inflammatory mediator. (F) Gene expression analysis using qPCR for IL6, a pro-inflammatory cytokine. Each cell line was differentiated into at least 3 clonal expansions, and the experiments were done in triplicates on those clonal lines. Data shown in bar graphs are presented as mean ± standard deviation and from at least three independent experiments. The p-value of <0.05 is considered significant. All confocal images shown are maximum projection of z-stack images. Scale bar: 50 microns

The extent of these changes differed between donor groups. AMD-derived cells exhibited more pronounced morphological alterations following treatment compared with non-AMD cells, with the most prominent changes observed in dry AMD-iPSC-RPE cells (Figure 3A).

The expression of genes associated with cellular stress and inflammatory responses was also examined. Treatment with CSE, PA, or the combination increased expression of *ZEB1*, *NQO1*, and *IL8* compared with untreated controls (Figure 3B-D). The increase was particularly pronounced in dry AMD-iPSC-RPE cells, which showed greater responses than non-AMD and advanced AMD-iPSC-RPE cells.

### CSE and PA alter RPE polarity and secretion of VEGF-A and PEDF

To examine the effect of CSE and PA on RPE polarity, mature iPSC-RPE cells were cultured on Transwell inserts and treated for 24 hours. Confocal imaging demonstrated disruption of the normal polarized distribution of ZO-1 and Best1 following CSE and PA exposure (Figure 4A). Quantification of the apical ZO-1 to basolateral Best1 fluorescence intensity ratio showed a significant reduction in polarity following treatment with CSE, PA, or the combined treatment in all cell lines (Figure 4B).

**Figure 4:**
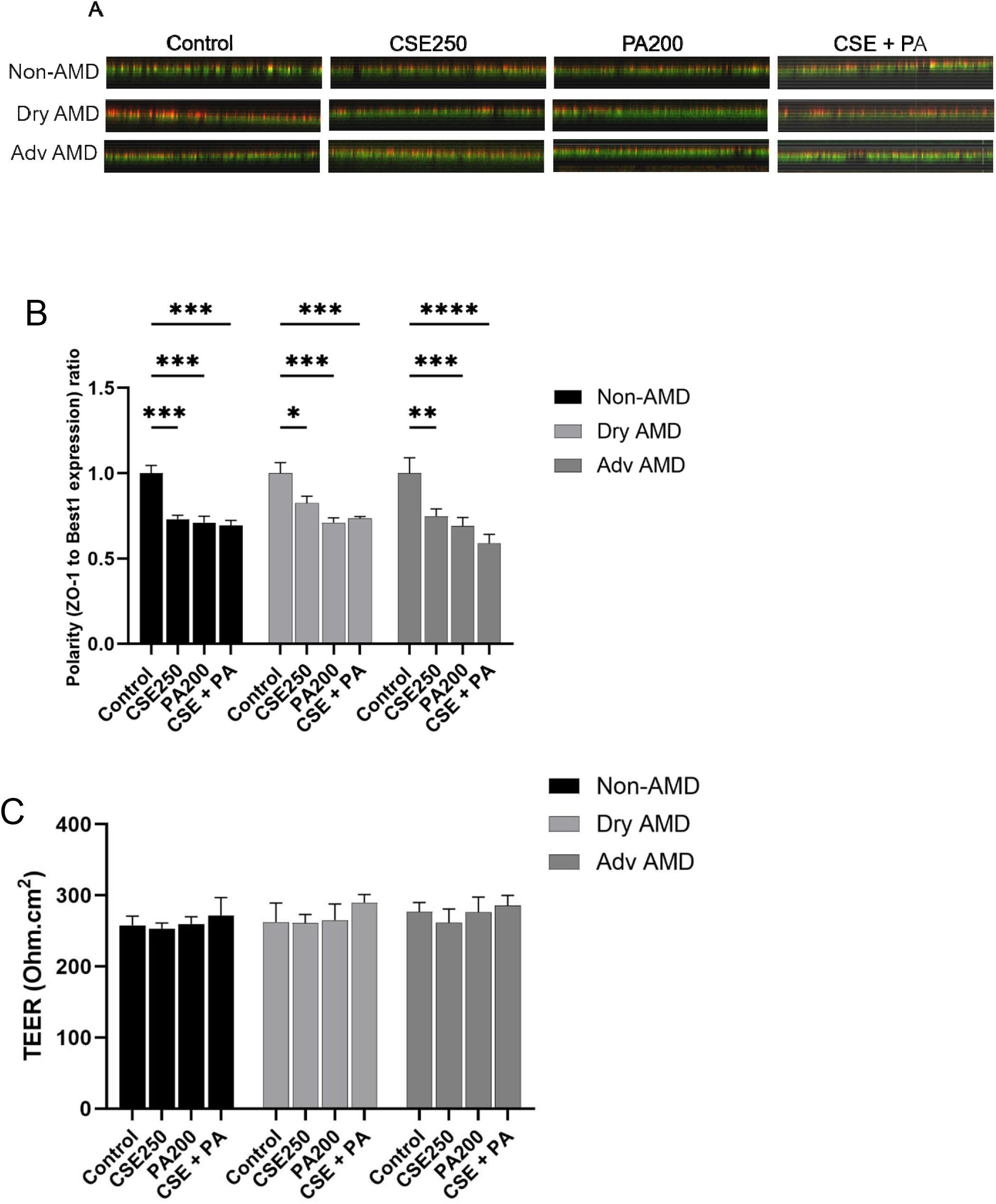
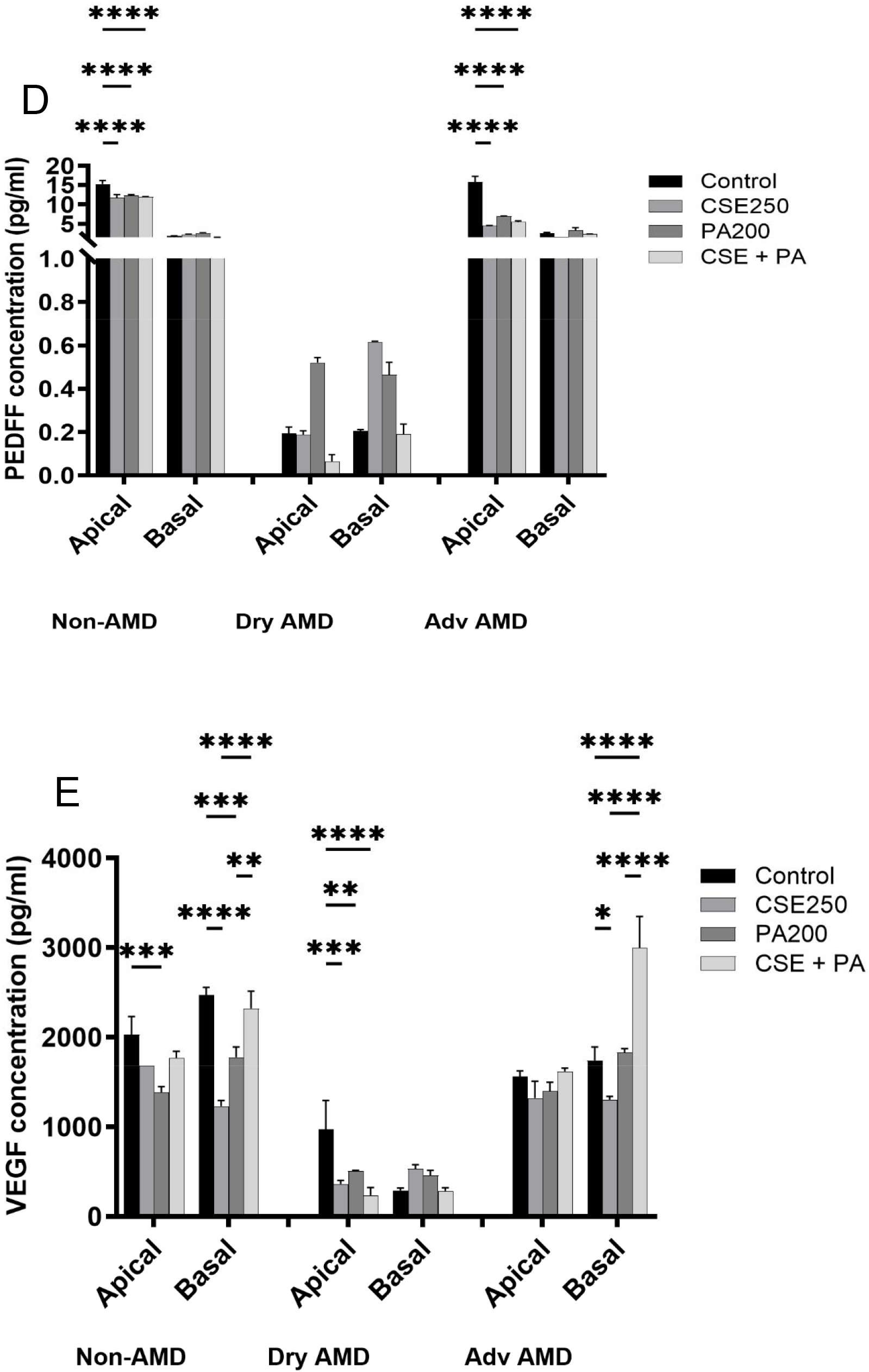
iPSC-RPE polarity post CSE and PA treatment. (A) Orthogonal confocal images showing ZO-1 and Best1 signals along the thickness of the iPSC-RPE monolayer. was used to identify the apical junctional plane. RPE polarity was quantified as the apical ZO-1 to basal Best1 fluorescence intensity ratio. (B) TEER measurement post treatment. (C) Quantification of ZO-1 and Best1 fluorescence intensity to measure polarity. ELISA analysis of human VEGF-A and PEDF secretion by non-AMD and AMD cell lines post treatments. Each cell line was differentiated into at least 3 clonal expansions, and the experiments were done in triplicates on those clonal lines. Data presented as bar graphs are representations of three independent experiments. *P < 0.05; **P < 0.01, ***P<0.001.

In contrast, TEER measurements did not show a significant difference between untreated and treated cells. TEER values remained above 200 Ω·cm² following treatment, with no significant differences observed between non-AMD and AMD-derived cells (Figure 4C).

The secretion of PEDF and VEGF-A was subsequently examined in the apical and basal compartments of the Transwell cultures. In non-AMD and Adv AMD-derived RPE cells, PEDF secretion was higher in the apical compartment than in the basal compartment. Treatment with CSE, PA, or their combination significantly reduced PEDF secretion compared with untreated controls in apical compartments. However, there were no changes in basal PEDF level with treatments for both cell lines. In contrast, dry AMD-iPSC-RPE cells exhibited a more balanced distribution of PEDF secretion between the apical and basal compartments and exposure to CSE or PA or both did not show any changes in VEGF levels in both apical and basal compartments (Figure 4D)

VEGF-A was detected in both apical and basal compartments, with higher secretion toward the basal side in mature iPSC-RPE cells. CSE and PA treatment significantly reduced VEGF-A secretion in both compartments when compared with untreated controls in non-AMD, while CSE and PA treatments significantly decreased VEGF-A secretion in apical compartment in dry AMD. In contrast, Adv AMD cells showed reduced VEGF secretion with CSE or PA treatment but increased significantly with combination treatment. . Dry AMD-iPSC-RPE cells exhibited lower VEGF-A secretion than the other two cell lines, with a more balanced distribution between the apical and basolateral compartments (Figure 4E).

### CSE and PA increase intracellular lipid accumulation

To determine whether CSE and PA affected lipid accumulation in iPSC-RPE cells, cells were treated with CSE, PA, or their combination for 24 hours and stained with BODIPY. Compared with untreated controls, CSE and PA exposure resulted in increased intracellular lipid accumulation, with BODIPY-positive lipid droplets observed within the cells (Figure 5A).

**Figure 5:**
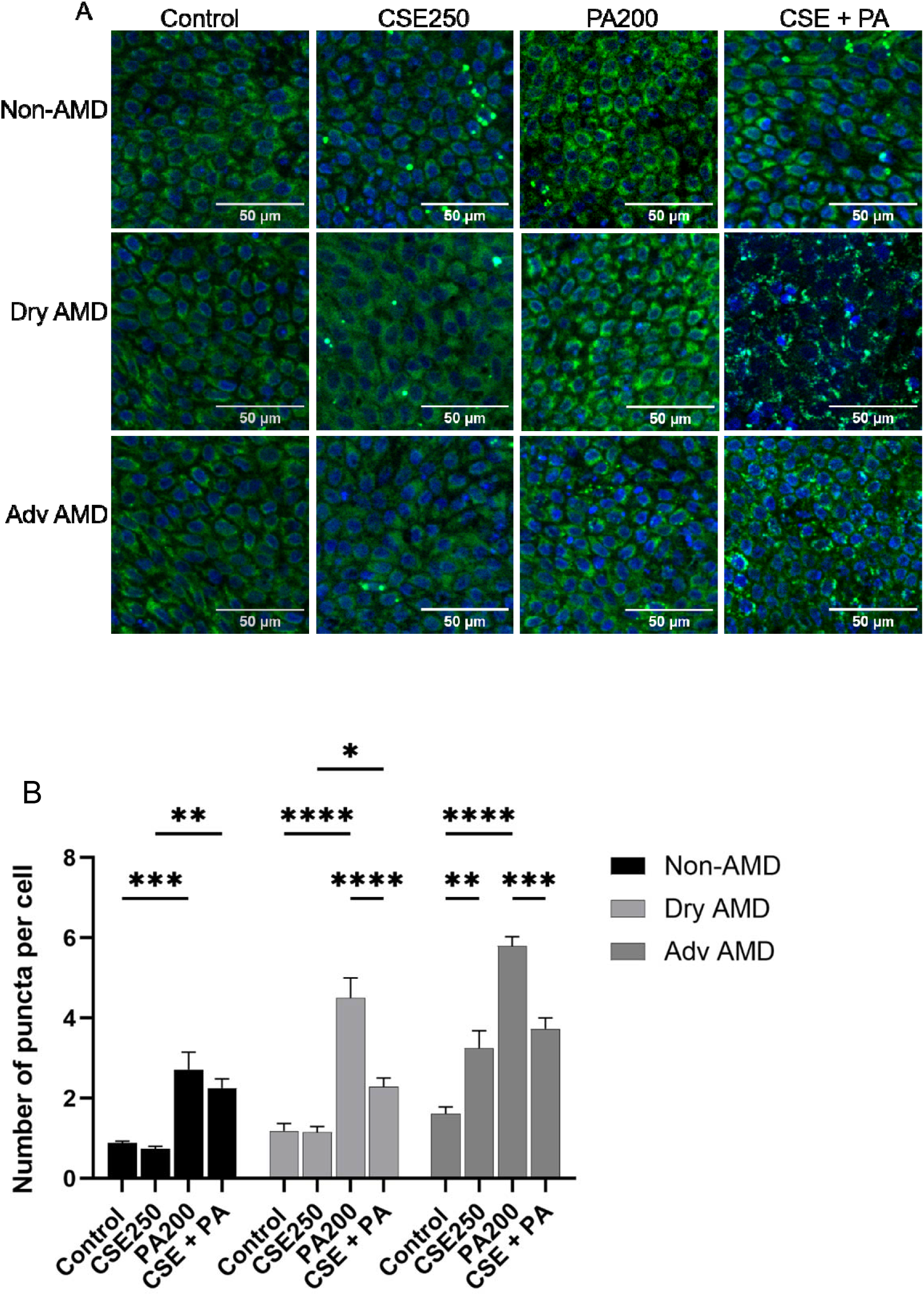
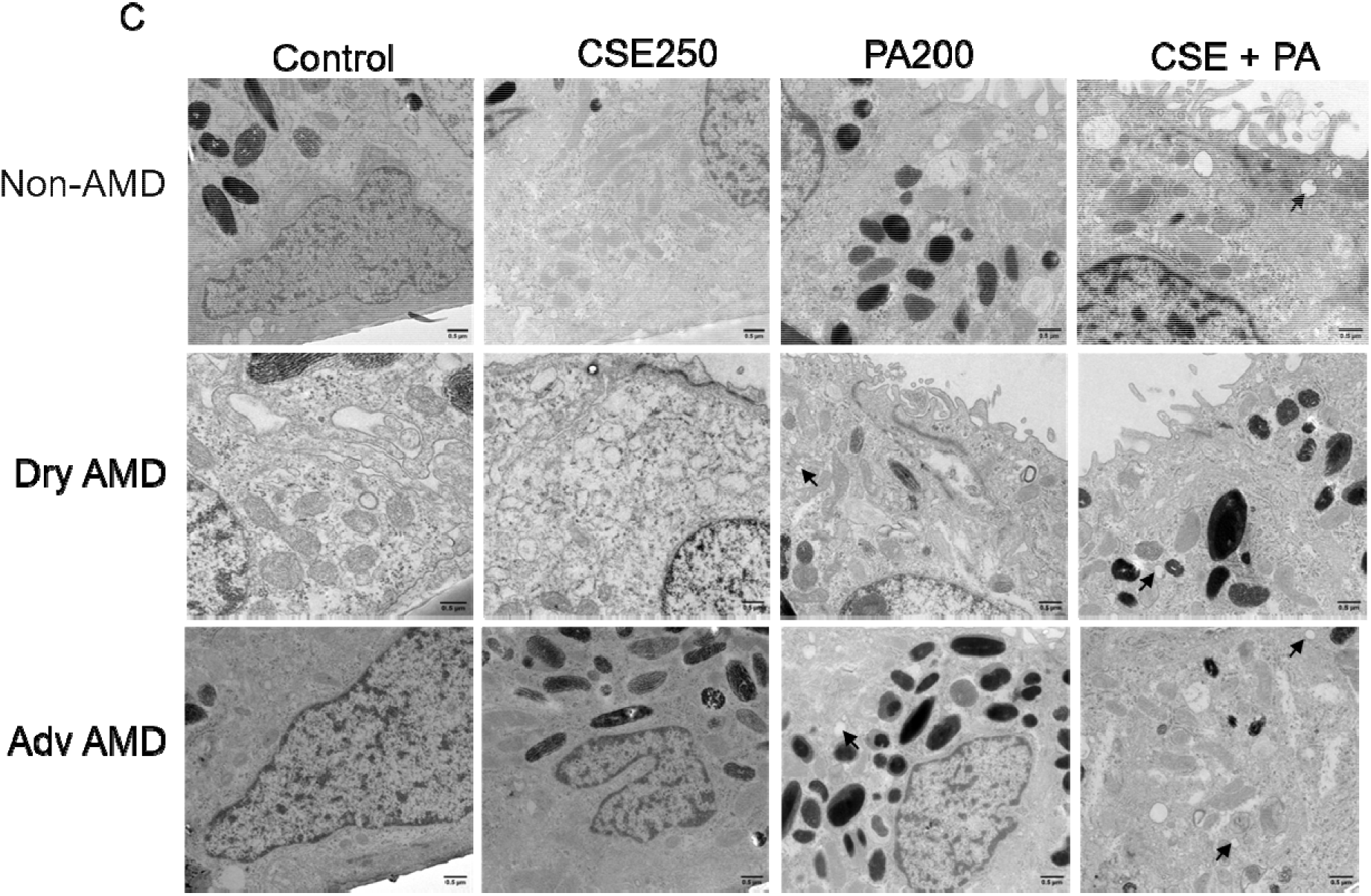
Lipid deposition in iPS-RPE cell monolayers exposed to cigarette smoke extract and PA. (A) Localization of lipids was identified using Bodipy stain. Bodipy stains particles were seen in RPE cells after CSE exposure compared to control. Lipid particles were increased with CSE exposure and PA exposure. (B) Quantification of Bodipy lipid particles after CSE and PA treatment for 24 hours. (C) TEM images showing lipid particles (indicated by black arrows) post CSE and PA treatments. Each cell line was differentiated into at least 3 clonal expansions, and the experiments were done in triplicates on those clonal lines. Data shown in bar graphs are presented as mean ± standard deviation and from at least three independent experiments. The p-value of <0.05 is considered significant. Images shown are representative images from the replicates.

Quantification of BODIPY staining showed a significant increase in the number of lipid particles per cell following PA treatment and combined CSE and PA treatment compared with untreated controls (Figure 5B). The increase in lipid accumulation was most pronounced in PA and combination treatment for all three cell lines. Despite the increase in intracellular lipid deposition, no corresponding significant change in TEER was observed.

Transmission electron microscopy further confirmed the presence of increased intracellular lipid droplets following PA and combined CSE and PA treatment (Figure 5C). These ultrastructural findings were consistent with the BODIPY staining results and further demonstrated increased lipid accumulation following exposure to metabolic and environmental stressors.

## Discussion

The use of patient-derived iPSC-RPE cells provides a human cell-based approach for investigating RPE responses relevant to AMD. The inclusion of cells derived from donors with clinically defined dry AMD and Adv AMD provides disease-relevant cellular backgrounds, while the non-AMD donor and the reference line provide comparative controls. In addition, the presence of AMD-associated genetic variants at loci including *CFH*, *ARMS2*, *C2*, and *C3* provides genetic context that may contribute to disease-associated cellular phenotypes. We specifically selected cell lines with a low-risk *CFH* genotype for the experiments with external-stressors to reduce the potential influence of *CFH*-associated genetic risk and allow the effects of the external stressors on iPSC-RPE cells to be examined under a more consistent *CFH* genetic background. This model therefore provides an opportunity to investigate how external stressors such as CSE and PA influence RPE phenotypes in the context of donor age and AMD disease status. However, given the complex multifactorial nature of AMD, which involves genetic, environmental, immune, and age-related factors, these iPSC-RPE models capture selected aspects of AMD pathophysiology rather than fully recapitulating the disease in vivo.

The study investigated the response of patient-derived iPSC-RPE cells to two AMD-relevant stressors: cigarette smoke extract (CSE), and palmitic acid (PA). The finding demonstrated that both treatments adversely affected iPSC-RPE cells, with AMD-iPSC-RPE cells generally exhibiting more pronounced responses than in non-AMD cells. In particular, dry AMD-iPSC-RPE cell line showed greater susceptibility to cell death, stress responses, changes in epithelial morphology, loss of polarity, and lipid accumulation. These findings suggest that AMD-derived RPE may have a reduced capacity to tolerate additional environmental and metabolic stress.

Before examining the effects of CSE and PA, we confirmed that the differentiated cells had characteristics of mature RPE. The cells developed pigmentation and the typical hexagonal morphology of RPE and expressed RPE-specific markers including *RPE65* and *TYRP1*. When cultured on Transwell inserts, the cells developed polarized localization of ZO-1 and Best1, with ZO-1 localized toward the apical region and Best1 toward the basolateral region. The TEER values were also above 200 Ω·cm² in all cell lines. Together with the downregulation of stem cell-associated markers and upregulation of RPE-specific genes, these findings indicate that the differentiated cells acquired several key structural and functional features of mature RPE. Importantly, AMD and non-AMD cells exhibited similar characteristics before treatment. This allowed us to examine differences in their response to stress rather than the potential influence of variations in RPE differentiation.

CSE and PA both produced a dose-dependent increase in cell death. The increased toxicity following CSE exposure is consistent with previous studies showing that cigarette smoke can induce oxidative stress and damage RPE cells. Smoking is one of the strongest modifiable environmental risk factors for AMD, and several mechanisms have been proposed, including oxidative damage, depletion of antioxidant defenses, inflammation, complement activation, and changes in lipid metabolism [6,11]. Our findings support that cigarette smoke exposure adversely affects RPE cells, with AMD iPSC-RPE cells potentially exhibiting greater susceptibility to its affects. PA also increased cell death, consistent with the effects of saturated fatty acid-induced lipotoxicity reported in other cell systems [19,20]. RPE cells have a high metabolic demand and are continuously exposed to lipids derived from photoreceptor outer segments. Maintaining lipid homeostasis is therefore important for RPE survival. An excess of saturated fatty acids may disturb this balance and lead to intracellular lipid accumulation and cellular stress. In the current study, AMD cells were more sensitive to PA than non-AMD cells, suggesting an altered capability to maintain lipid and metabolic homeostasis.

Combined treatment with CSE and PA resulted in greater cell death than either treatment alone. This finding is particularly relevant to AMD, as environmental and metabolic risk factors are likely to act together over long periods rather than independently. Previous studies have demonstrated that cigarette smoke exposure induces oxidative stress, complement activation, endoplasmic reticulum stress, and lipid accumulation in RPE cells [9,21]. Therefore, the stronger response observed with combined CSE and PA treatment may reflect the ability of these two stressors to affect overlapping cellular pathways. Although our findings do not establish a specific synergistic mechanism, they suggest that simultaneous exposure to environmental and lipid stressors exacerbates RPE cell vulnerability.

An important observation in this study was that AMD-iPSC-RPE cells were more susceptible to CSE and PA-induced cell death than non-AMD cells. This difference was particularly apparent in dry AMD-derived cells. RPE dysfunction and loss are central features of non-neovascular AMD, and oxidative stress and inflammation are thought to contribute to the progression of RPE degeneration [22,23]. The greater sensitivity of dry AMD-derived RPE in our model may therefore indicate an intrinsic difference in the ability of these cells to respond to additional stress. The mechanism underlying this susceptibility remains unclear. Differences in antioxidant defense capacity, mitochondrial function, lipid metabolism, or inflammatory signaling may potentially contribute to this vulnerability and require further investigation.

CSE and PA also produced morphological changes consistent with an EMT-like response. The changes were more pronounced in AMD cells, particularly in the dry AMD group. The increase in *ZEB1* expression following treatment provides additional support for an alteration in the epithelial phenotype. RPE cells normally maintain a highly differentiated epithelial phenotype, and disruption of this phenotype can affect cell-cell interactions, polarity, barrier properties, and other functions required for retinal homeostasis. Although the present study did not examine a complete panel of EMT markers, the morphological changes together with increased *ZEB1* expression suggest that CSE and PA can promote an EMT-like state in RPE cells. Further studies examining additional epithelial and mesenchymal markers would be needed to determine the extent of this transition.

The increase in *NRF2*, *NQO*, *IL6* and *IL8* expressions, further suggests that CSE and PA activate cellular stress pathways. *NRF2* and *NQO1* are part of the cellular antioxidant and detoxification response and is regulated by pathways involved in protection against oxidative stress. Cigarette smoke is known to generate oxidative stress in the RPE and can activate antioxidant responses [11]. The increase in *NRF2* and *NQO1* observed here may therefore represent a protective response to CSE- and PA-induced stress. In contrast, the increase in *IL6* and *IL8* suggests activation of an inflammatory response after CSE and PA treatment. Inflammation has an important role in AMD, and chronic oxidative and inflammatory stress may contribute to progressive RPE dysfunction [22]. The stronger response in dry AMD cells for these stress response genes again suggests that these cells may have a lower threshold for activation of stress and inflammatory pathways.

Another significant finding was the effect of CSE and PA on RPE polarity. Under control conditions, ZO-1 and Best1 showed the expected polarized distribution. Following treatment, this distribution was disrupted, and the ratio of apical ZO-1 to basolateral Best1 fluorescence was significantly reduced. RPE polarity is important for maintaining normal retinal function because the apical and basolateral surfaces of the RPE have distinct roles in transport, communication with photoreceptors, and interaction with the choroidal circulation. Polarization also influences the secretion of important RPE-derived factors. Previous studies have shown that polarized RPE preferentially secretes PEDF toward the apical side and VEGF toward the basolateral side [24].

Interestingly, TEER remained unchanged following CSE or PA treatment despite alterations in ZO-1 and BEST1 localization. These findings suggest that changes in RPE epithelial organization can occur without a measurable reduction in barrier resistance. Although TEER is a useful indicator of epithelial barrier integrity, it does not fully reflect changes in tight junction organization or apicobasal polarity. Therefore, the absence of significant changes in TEER does not necessarily indicate that RPE function remains unaffected. Our findings suggest that CSE and PA can alter RPE epithelial organization while barrier resistance remains relatively preserved.

Changes in PEDF and VEGF secretion provide additional evidence of functional alterations following treatment. PEDF was preferentially secreted toward the apical side in the non-AMD and Adv AMD cells under control conditions, which is consistent with previous studies of polarized RPE [24,25]. CSE and PA significantly reduced PEDF secretion. This reduction may be relevant because PEDF has neuroprotective and anti-angiogenic properties and is an important component of the RPE secretome. The altered distribution of PEDF in dry AMD cells, characterized by a more balanced between the apical and basal compartments, may also indicate an underlying alteration in RPE polarity or secretory function. In contrast, VEGF-A was detected in both compartments but exhibited preferential basolateral secretion, consistent with previous reports in differentiated RPE [26,27]. Basolateral VEGFA plays an important physiological role in supporting choriocapillaris and maintaining the RPE-choroidal interface. Consequently, the reduction in VEGF secretion observed after CSE and PA treatment may reflect impaired RPE rather than merely a reduction in angiogenic activity. Furthermore, the lower VEGF secretion observed in dry AMD-derived cells suggests potential differences in the baseline secretory phenotype of these cells.

Increase in intracellular lipid accumulation following PA and CSE exposure was another important finding. BODIPY staining revealed increased lipid deposition following PA treatment and combined CSE and PA treatment, and this was supported by transmission electron microscopy. The higher level of lipid accumulation in dry AMD cells is particularly interesting given the importance of lipid dysregulation in AMD pathogenesis. Alteration in lipid accumulation and changes in lipid composition are associated with aging and with extracellular deposits in the RPE-Bruch’s membrane complex. Previous studies have also shown that cigarette smoke exposure can induce lipid accumulation in RPE cells through oxidative stress, complement activation, and endoplasmic reticulum stress [9,28]. Together, our findings suggest that cigarette smoke exposure and excess saturated fatty acid can independently or jointly disturb lipid homeostasis in RPE.

The relationship between lipid accumulation and cell death is also important to consider. Lipid droplets can initially serve as a means of storing excess fatty acids and may protect cells from the accumulation of toxic free fatty acids. However, excessive lipid accumulation may contribute to oxidative stress, mitochondrial dysfunction, inflammation, and ultimately cell death. The greater lipid accumulation observed in dry AMD-derived cells could therefore reflect either increased lipid uptake/storage or a reduced capacity to metabolize and clear excess fatty acids. Further investigation of lipid composition, fatty acid oxidation, and lipid storage and degradation pathways would help clarify the mechanisms underlying these alterations.

Overall, the findings suggest that CSE and PA affect several aspects of RPE biology. Both stressors increased cell death and induced changes in cellular morphology and stress-related gene expression. They also disrupted RPE polarity, altered the secretion of PEDF and VEGF, and increased intracellular lipid accumulation. These effects were generally more pronounced in AMD-derived cells, with dry AMD cells showing the greatest susceptibility. The more pronounced effects observed following combined CSE and PA treatment further suggest that environmental and metabolic stressors may act together to exacerbate RPE dysfunction.

The differences between dry AMD and Adv AMD cells are particularly interesting. Although both groups were derived from AMD patients, the dry AMD cells consistently showed greater sensitivity to several of the treatments. These differences may reflect variations in disease stage, donor-specific factors, or intrinsic differences in cellular stress responses. Because iPSC-derived RPE cells retain certain donor-specific characteristics, this model may provide a valuable approach for investigating the heterogeneity of AMD. However, the relatively limited number of donor-derived cell lines should be considered when interpreting these differences, and additional donors will be necessary to determine whether the observed phenotype is consistently associated with dry AMD.

There are several limitations to this study. First, the RPE does not function independently *in-vivo*, and the iPSC-RPE model does not reproduce the complete retinal environment, including interactions with photoreceptors, choroidal endothelial cells, immune cells, and Bruch’s membrane. Second, the 24-hour exposure used in this study represents an acute stress condition, whereas smoking and metabolic abnormalities occur chronically in patients. Similarly, PA represents only one type of fatty acid and does not fully reproduce the lipid environment associated with aging and AMD. Finally, additional functional assays will be needed to determine whether the observed changes in polarity, lipid accumulation, and secretion directly contribute to RPE cell dysfunction. Despite these limitations, the present study suggests that AMD-derived RPE cells respond differently to environmental and metabolic stress than non-AMD RPE cells. The increased susceptibility of dry AMD-derived cells to CSE and PA is particularly relevant because RPE dysfunction and degeneration are major features of the disease. The findings also support the use of patient-derived iPSC-RPE as a model for studying interactions between genetic or disease-associated susceptibility and environmental risk factors.

In conclusion, our data suggests that cigarette smoke exposure and excess palmitic acid can independently and jointly disrupt RPE homeostasis. These stressors induced cell death, EMT-like changes, cellular stress and inflammatory responses, alteration in RPE polarity and growth factor secretion, and increased intracellular lipid accumulation. The stronger response observed in AMD-derived cells, particularly dry AMD cells, suggests that disease-associated RPE may be less capable of adapting to additional environmental and metabolic stress. Further investigation of the molecular pathways underlying this increased susceptibility may provide insights into how modifiable risk factors, such as smoking and altered lipid metabolism, contribute to RPE dysfunction and AMD progression.

## Material and Methods

### Cell Lines Used

Fibroblast derived iPSC lines from one normal (non-AMD), one dry AMD and one geographic atrophy (GA) individuals were obtained from Golestaneh lab at Georgetown University, Washington and IMR90-4 was obtained from WiCell (University of Wisconsin–Madison, USA). Demographic data (age, sex) and disease status were collected for the first three lines. Disease status was determined by qualified clinicians. Genotyping for key AMD related pathways was performed as detailed below.

### Genotyping

Genomic DNA was extracted from mature iPS-RPE cells using All prep DNA/RNA Mini kit (Qiagen; Hilden, Germany). DNA was quantified using Agilent BioTek Take3 microvolume plate using 2µl of undiluted genomic DNA sample in an Agilent BioTek SYNERGY H1 microplate reader (Biotek, USA). Samples were genotyped for the *CFH* (rs1061170), *ARMS2* (rs10490924), *C2* (rs547154) and *C3* (rs2230199) using allele-specific taqman primers designed for the single nucleotide polymorphism (SNP) (Thermo Fisher Scientific).

### Cell culture and maintenance

The frozen iPSCs were thawed in culture flasks coated with human recombinant vitronectin (0.5ug/ml, Life technologies A14701SA) diluted in DPBS without CaCl_2_ and MgCl_2._ The vitronectin coated plates were incubated at 37°C overnight and the liquid was removed before use. For thawing media, Rock Inhibitor (Y-27632) was added to E8 complete media to achieve final concentration of 10µM. The cells were fed with E8 complete media daily until the culture reached 70-80% confluency or ready to be passaged or harvested. Differentiation of iPSCs to mature RPE cells were done using previously described protocol [29]. The cells were allowed to differentiate for 10 weeks to obtain mature iPS-RPE cells. Briefly, the confluent iPSCs colonies were dissociated with Versene (Life technologies, 15040066) and seeded to new wells coated with vitronectin using plating media (Essential 8 complete media containing 10µg/ml blebbistatin). Cells were maintained in E8 complete media for two days with daily media replacement. The iPSCs were then switched to Retinal Induction Medium (RIM) for two days and cultured as Neuroectoderm cells with medium changed every day. Then the culture medium was switched to Retinal Differentiation Media (RDM) and continued for 9 days feeding every day as RPE progenitor cells. The RPE progenitor cells can be seen some neuronal clusters which can be removed by passing the dissociated cells through 20µm steriflip cell strainer. Filtered cells were fed with Retinal Media with Nicotinamide and Activin-A (RM-NA) every day, for 11 days and cultured as committed RPE cells. RPE cells were then fed with RPE Maturation Media (RPEMM) every other day and cultured as immature RPE cells and kept growing until maturation. The immature RPE cells can be dissociated and seeded to multiple vitronectin coated wells or Transwells for maturation in RPE plating media. The medium was changed to RPE maturation media (RPEMM with Prostaglandin 2) at 10 days post re-seeding, to enhance maturation of iPS-RPE cells. For Transwells, the media were replenished in top and bottom compartments separately. Each cell line was differentiated into at least 3 clonal expansions, and the experiments were done on those clonal lines in triplicates.

### CSE and PA treatment

The cigarette smoke extract used in treatments was purchased from Murty Pharmaceuticals as a 40 mg/ml stock. Working solutions for CSE were prepared by diluting the stock in 100% DMSO. The PA used in treatments was conjugated to fatty-acid-free BSA as described previously. Briefly, PA was dissolved in anhydrous ethanol in a water bath at 50 °C to yield a final concentration of 25 mM. In parallel, BSA was dissolved in water to get a final concentration of 0.5 mM. After its solubilization, PA was mixed with BSA to obtain a 6:1 molar ratio. The solution was then vortexed and incubated at 55 °C for PA-BSA conjugation to occur. The PA-BSA mix was cooled to room temperature, filter sterilized and stored at -20 °C until use. For each experiment, DMSO only control for CSE treatment and BSA only control for PA treatment were included.

### Cytotoxicity assay

To assess the impact of CSE and PA on cell survival, cells were treated with CSE or PA or both and stained with Ethidium homodimer-1 (Thermo Fisher, E1169) to detect dead or dying cells. Hoechst stain was used for total cells nuclear stain. Briefly, the cells were cultured on a 96 well plate to achieve 70-80% confluency and treated with CSE and PA for 24 hours. The staining solution was prepared by adding 20 µl ethidium homodimer-1 (Invitrogen, E1169) to 10 ml DPBS. The culture media was removed from the cells, and the 100-200 µl of the staining solution was added directly to the cells. The cells were incubated for 30 minutes at 37°C and washed with PBS. Then the cells were counter stained with 1 µg/ml Hoechst for 10 mins and washed with PBS. The imaging buffer was added to the cells and images were captured using an EVOS microscope.

### RNA isolation, cDNA synthesis and qPCR

Total RNA was extracted from patient-derived iPS-RPE cells using the Quick-RNA^TM^ MiniPrep Kit (Zymo Research Corporation R1055). Complementary DNA (cDNA) was synthesized using the High-Capacity cDNA Synthesis kit (Thermo Fisher) using C1000 Touch Thermal Cycler (BioRad). qPCR for individual gene expression was performed using predesigned Taqman primer probes (Thermo Fisher) and the reactions were carried out on the CFX Opus 96 Real Time PCR system (Bio-Rad). Gene expressions were quantified by the relative quantification method of 2^−ΔΔCt^ and normalized to housekeeping genes.

### Immunocytochemistry and confocal imaging

Cells were fixed in 4% paraformaldehyde in phosphate-buffered saline (PBS) for 15 min at room temperature (RT) followed by permeabilization using 0.1% (vol/vol) Triton X-100 in PBS for 30 minutes and blocking in 1% (wt/vol) Bovine Serum Albumin (BSA) in PBST (0.01% Tween20 in 1X PBS) with 0.3M glycine for 1h at RT. For transwells, the reagents were added separately to the top and bottom of the chamber. Primary antibodies used in this study were anti-ZO1 conjugated with Alexa 594 (Thermo Fisher, 187430), anti-Best1 (Novus Biologicals), anti-TYRP1 (Novus Biologicals, NBP2-32907), anti-REP65 (From Micah). The slides were mounted with Vectashield vibrance with DAPI to counter stain nuclei. Images were captured using a Leica-Mica confocal microscope. Images were captured as z-stacks with 1µm step size.

### Transepithelial electrical resistance (TEER) measurements

Transepithelial resistance (TER) was measured for iPSC-RPE cells cultured on vitronectin-coated 12 well-transwell inserts, using an EVOM2 epithelial voltohmmeter with a STX2 chopstick electrode (World Precision Instruments Inc., USA). Briefly, the electrode was sterilized using 70% ethanol and equilibrated into pre-warmed culture medium, before being introduced into apical and basal transwell compartments. Measurements were recorded for at least three separate wells per cell line per experiment. Fo each well, at least three readings were taken to obtain an average value. The net TEER was determined by subtracting the resistance across a vitronectin-coated transwell insert lacking cells, from measured values, and then multiplying the growth surface area of the wells as shown in equation1 below.

Final TEER (Ω/cm^2^) = Net TEER (Ω) × Area of Transwell insert (cm^2^)

### VEGF and PEDF secretion assays

Spent medium was collected from both compartments of transwell inserts for further analysis of secreted factors using commercially available enzyme-linked immunosorbent assay kits for human Vascular Endothelial Growth Factor-A (VEGF-A) and human Pigment Epithelium-Derived Factor (PEDF). To determine secretion properties, iPSC-RPE monolayers were grown for 10 weeks on vitronectin-coated 12-well transwell inserts. Apical and basal VEGF-A and PEDF levels were measured using the commercially available human VEGF ELISA Kit (Sigma, RAB0507) and human PEDF Chemikine ELISA Kit (Sigma, CYT420) following the manufacturer’s instructions. Microplate reading was performed at 450nm with the SYNERGY H1 microplate reader (Biotek).

### RPE polarity

The mature cells attached to the transwell membrane were processed for immunocytochemistry and stained with ZO-1 and Best1 as apical and basal markers respectively. Orthogonal confocal images were analyzed in ImageJ. The ZO-1 signal was used to identify the apical plane, and *BEST1* signal was used to identify basal planes. A rectangular region of interest (ROI) encompassing the full thickness of the RPE monolayer was divided into apical and basal compartments. The mean fluorescence intensity of ZO-1 and BEST1 was measured in apical and basal compartment respectively after background subtraction. RPE polarity was quantified as the apical ZO-1-to-basal BEST1 fluorescence intensity ratio. A ratio higher than 1 indicates a highly polarized monolayer where tight junctions are sharply defined at apical end relative to the basolateral region. A ratio of less than 1 suggests loss of epithelial polarity potentially.

### Protein extraction and Immunoblot

For protein extraction, the cell pellets were added with RIPA buffer (Thermo Fisher Scientific) containing protease and phosphatase inhibitor cocktail (Thermo Fisher Scientific, 78440). Lysates were rotated using a multi-tube rotator for 20 minutes at 4°C, followed by centrifugation at 10,000g for 5 minutes at 4°C. The supernatant was collected, and the protein concentration was estimated using DC assay. Equal amount of protein samples was mixed with Laemmli sample buffer containing 5% β-mercaptoethanol and boiled at 95°C for 5 minutes. Proteins were separated on 4-12% NuPAGE^TM^ Bis-Tris mini gels (Life technologies, NP0336BOX) and transferred to 0.2µm nitrocellulose membrane (Bio-Rad, Trans-Blot Turbo Mini 0.2µm Nitrocellulose transfer Packs, 1704158) using a turbo transfer system (Bio-Rad). The proteins on the membrane were blocked in 5% non-fat dry milk in TBS-T (Tris-buffered saline with 0.1% Tween-20) or 5% Bovine Serum Albumin (BSA) for 1h at RT on a shaker. The blots were washed 4 times with 1X TBS-T and incubated overnight at 4°C with primary antibodies. After washing with TBS-T, membranes were incubated with HRP-conjugated secondary antibodies (anti-rabbit or anti-mouse) for 1 hour at room temperature and washed with TBS-T. For loading control, the washed blot was incubated with HRP-conjugated anti-beta actin antibody for 1h at RT. Detection was performed using enhanced chemiluminescence (ECL) reagents (SuperSignal West, Thermo Fisher Scientific) and imaged using a chemiluminescence detection system (Bio-Rad).

### Transmission electron microscopy (TEM)

Cells grown on transwell inserts were processed for transmission electron microscopy (TEM). Briefly, the transwell membranes with cells were washed in 1X PBS, followed by immersion in primary fixative (2.5% glutaraldehyde in 0.1 M cacodylate buffer; pH 7.4) at 4°C overnight. The cells were rinsed twice for 10 minutes in 0.1M PIPES and post fixed in 1% osmium tetroxide in 0.1M PIPES for 1h at room temperature. Transwell inserts were rinsed twice in 0.1M PIPES and once in ddH2O, and en bloc stained in 2% aqueous uranyl acetate for 20 minutes. Samples were then subsequently dehydrated by treating through a graded ethanol concentration series (30%, 50%, 70% and 95%) for 10 minutes each, followed by two 20-minute incubations in absolute ethanol (100%). The membranes were then embedded in epoxy resin. Ultrathin sections were cut using an ultramicrotome and collected on collected on 200 mesh carbon and formvar coated copper grids and were counterstained with uranyl acetate and lead citrate prior to imaging Cross sections of cells on transwell inserts were visualized using a transmission electron microscope operated at 80-120 kV. Images were acquired to assess the ultrastructure of the retinal pigment epithelium (RPE) cells with particular attention to mitochondrial morphology and sub-RPE basal laminar deposits. Representative images were captured at multiple magnifications.

### Statistical analysis

Data are presented as mean ± standard deviation of at least three independent experiments. Comparisons between two or more treatments were performed using two-way analysis of variance followed by Tukey’s post hoc test to correct for multiple comparisons. A p-value <0.05 was considered statistically significant. Statistical analyses were performed using GraphPad Prism for Windows version 8, (GraphPad Software, San Diego, CA, USA).

