## Supplementary Material for "Cigarette smoke extract and palmitic acid exposed patient-derived iPS-RPE cells as a model for AMD"

**Supplementary Data**

Supplementary Table 1: SNPs genotyping result for iPSC-RPE lines

| iPSC-RPE line | ^a^Age/^b^Gender | Genotype | | | | | | | |
| --- | --- | --- | --- | --- | --- | --- | --- | --- | --- |
|  |  | ^c^*CFH* | Risk | ^d^*ARMS2* | Risk | ^e^*C2* | Risk | ^f^*C3* | Risk |
| IMR90-4 (Reference line) | N/A | CT | Moderate | GG | Low | GT | Low | GG | High |
| Non-AMD (family history of AMD) | 75/F | TT | Low | GT | Moderate | GT | Low | GG | High |
| Dry AMD | 86/M | TT | Low | TT | High | GT | Low | CG | Moderate |
| Advanced stage AMD (Adv AMD) | 80/F | TT | Low | TT | High | GT | Low | CG | Moderate |

^a^Age of donor patients in years. ^b^Gender of donor patients. ^c^Complement facor H (*CFH*) genotype for rs1061170; low risk = TT, moderate risk = CT, high risk = CC. ^d^Age Related Maculopathy Susceptibility 2 (*ARMS2*) genotype for rs10490924; low risk = GG, moderate risk = GT, high risk = TT. ^e^Complement component 2 (*C2*) genotyping for rs547154; low risk = GT or TT, high risk = GG. ^f^Complement component 3 (*C3*) genotyping for rs2230199; low risk = CC moderate risk = CG, high risk = GG.

Supplementary Figure1 Western blot analysis of Nrf2 expression across CSE and PA treated non-AMD and AMD iPS-RPE cells

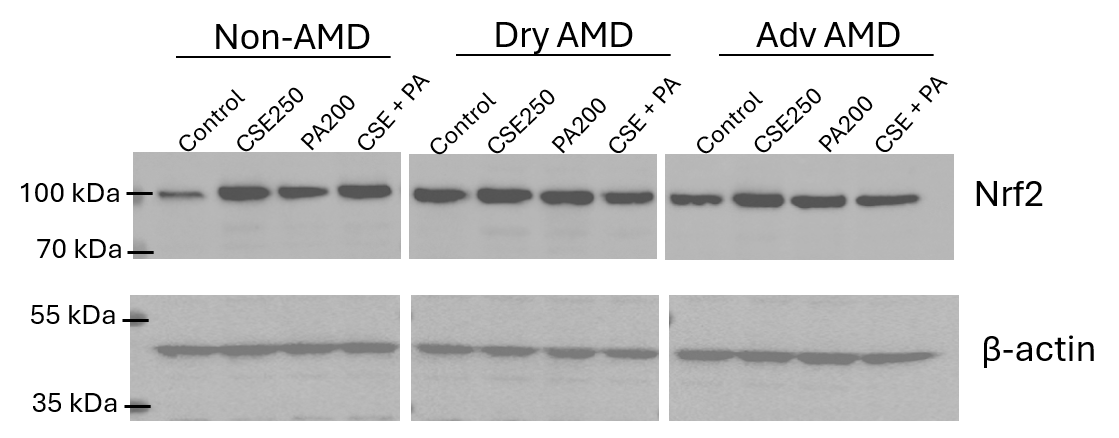

Representative immunoblots showing the expression levels of Nrf2 (top panel) and β-actin (bottom panel) following CSE and PA treatments. Samples are organized into three groups representing three cell lines: Non-AMD, Dry AMD and Adv AMD with four treatment conditions per cell line. The relatively uniform signal intensity of the loading control across lanes confirms equal protein loading. Molecular weight markers are indicated on the left of each panel. Each cell line was differentiated into at least 3 clonal expansions, and the experiments were done in triplicates on those clonal lines.
